# Deep learning and computer vision will transform entomology

**DOI:** 10.1101/2020.07.03.187252

**Authors:** Toke T. Høye, Johanna Ärje, Kim Bjerge, Oskar L. P. Hansen, Alexandros Iosifidis, Florian Leese, Hjalte M. R. Mann, Kristian Meissner, Claus Melvad, Jenni Raitoharju

## Abstract

Most animal species on Earth are insects, and recent reports suggest that their abundance is in drastic decline. Although these reports come from a wide range of insect taxa and regions, the evidence to assess the extent of the phenomenon is still sparse. Insect populations are challenging to study and most monitoring methods are labour intensive and inefficient. Advances in computer vision and deep learning provide potential new solutions to this global challenge. Cameras and other sensors that can effectively, continuously, and non-invasively perform entomological observations throughout diurnal and seasonal cycles. The physical appearance of specimens can also be captured by automated imaging in the lab. When trained on these data, deep learning models can provide estimates of insect abundance, biomass, and diversity. Further, deep learning models can quantify variation in phenotypic traits, behaviour, and interactions. Here, we connect recent developments in deep learning and computer vision to the urgent demand for more cost-efficient monitoring of insects and other invertebrates. We present examples of sensor-based monitoring of insects. We show how deep learning tools can be applied to the big data outputs to derive ecological information and discuss the challenges that lie ahead for the implementation of such solutions in entomology. We identify four focal areas, which will facilitate this transformation: 1) Validation of image-based taxonomic identification, 2) generation of sufficient training data, 3) development of public, curated reference databases, and 4) solutions to integrate deep learning and molecular tools.

**Significance statement:** Insect populations are challenging to study, but computer vision and deep learning provide opportunities for continuous and non-invasive monitoring of biodiversity around the clock and over entire seasons. These tools can also facilitate the processing of samples in a laboratory setting. Automated imaging in particular can provide an effective way of identifying and counting specimens to measure abundance. We present examples of sensors and devices of relevance to entomology and show how deep learning tools can convert the big data streams into ecological information. We discuss the challenges that lie ahead and identify four focal areas to make deep learning and computer vision game changers for entomology.

## INTRODUCTION

We are experiencing a mass extinction of species (1), but data on changes in species diversity and abundance have substantial taxonomic, spatial, and temporal biases and gaps (2, 3). The lack of data holds especially true for insects despite the fact that they represent the vast majority of animal species. A major reason for these shortfalls for insects and other invertebrates is that available methods to study and monitor species and their population trends are antiquated and inefficient (4). Nevertheless, some recent studies have demonstrated alarming rates of insect diversity and abundance loss (5-7). To further explore the extent and causes of these changes, we need efficient, rigorous, and reliable methods to study and monitor insects (4, 8).

Data to derive insect population trends are already generated as part of ongoing biomonitoring programs. However, legislative terrestrial biomonitoring, e.g. in the context of the EU Habitat Directive, focuses on a very small subset of individual insect species such as rare butterflies and beetles because the majority of insect taxa are too difficult or too costly to monitor (9). In current legislative aquatic monitoring, benthic invertebrates are commonly used in assessments of ecological status (e.g. the US Clean Water Act, the EU Water Framework Directive, and the EU Marine Strategy Framework Directive). Still, the spatiotemporal and taxonomic extent and resolution in ongoing biomonitoring programs is coarse and does not provide information on the status of the vast majority of insect populations.

Molecular techniques such as DNA barcoding and metabarcoding will likely become valuable tools for future insect monitoring based on field collected samples (10, 11), but at the moment high-throughput methods cannot provide reliable abundance estimates (12, 13) leaving a critical need for other methodological approaches. The state-of-the-art in deep learning and computer vision methods and image processing has matured to the point where it can aid or even replace manual observation *in situ* (14) as well as in routine laboratory sample processing tasks (15). Image-based observational methods for monitoring of vertebrates using camera traps have undergone rapid development in the past decade (14, 16-18). Similar approaches using cameras and other sensors for investigating diversity and abundance of insects are underway (19, 20). However, despite huge attention in other domains, deep learning is only very slowly beginning to be applied in invertebrate monitoring and biodiversity research (21-25).

Deep learning models learn features of a dataset by iteratively training on example data without the need for manual feature extraction (26). In this way, deep learning is qualitatively different from traditional statistical approaches to prediction (27). Deep learning models specifically designed for dealing with images, so called convolutional neural networks (CNNs) can extract the features of various aspects of a set of images or the objects within them, and learn to differentiate among them. There is great potential in automatic detection and classification of insects in video or time-lapse images with trained CNNs (20). As the methods become more refined, they will bring exciting new opportunities for understanding insect ecology and for monitoring (19, 28-31).

Here, we argue that deep learning and computer vision can be used to develop novel, high throughput systems for detection, enumeration, classification, and discovery of species as well as for deriving functional traits such as biomass for biomonitoring purposes. These approaches can help solve long standing challenges in ecology and biodiversity research and the pressing issues in insect population monitoring (32, 33). This article has three goals. First, we present sensor-based solutions for observation of invertebrates *in situ* and for specimen-based research in the laboratory, which due to the volume of data generated, use or could benefit from deep learning models to process data. Second, we show how deep learning models can be applied to obtained data streams to derive ecologically relevant information. Last, we outline and discuss four main challenges that lie ahead in the implementation of such solutions for invertebrate monitoring, ecology, and biodiversity research.

## SENSOR-BASED INSECT MONITORING

Sensors are widely used in ecology for gathering peripheral data such as temperature, precipitation, light intensity etc., but have not yet been used much for gathering data on the insects. However, solutions for sensor-based monitoring of insects and other invertebrates in their natural environment are emerging (34). The innovation and development is primarily driven by agricultural research to predict occurrence and abundance of beneficial and pest insect species of economic importance (35-37), to provide more efficient screening of natural products for invasive insect species (38), or to monitor disease vectors such as mosquitos (39, 40). The most commonly used sensors are cameras, radar, and microphones. Such sensor-based monitoring is likely to generate big data, which require efficient solutions for extracting relevant biological information. Deep learning could be a critical tool in this respect. Below, we give examples of image-based approaches to insect monitoring, which we argue has the greatest potential for integration with deep learning. We also describe approaches using other types of sensors, where the integration with deep learning is less well developed, but still could be relevant for detecting and classifying entomological information. We further describe the ongoing efforts in the digitization of natural history collections, which could generate valuable reference data for training and validating deep learning models.

### Image-based solutions for *in situ* monitoring

Some case studies have already used cameras and deep learning methods for detecting single species, such as the pest of the fruits of olive trees *Bactrocera oleae* (41) or for more generic pest detection (42). The pest detection is based on images of insects that have been trapped with either a McPhail-type trap or a trap with pheromone lure and adhesive liner. The images are collected by a microcomputer and transmitted to a remote server where they are analysed. Other solutions have embedded a digital camera and a microprocessor that can count trapped individuals in real-time using object-detection based on an optimized deep learning model (37). In both these cases, deep learning networks are trained to recognize and count the number of single pest species. However, there are very few examples of invertebrate biodiversity-related field studies applying deep learning models (23). Early attempts used feature vectors extracted from single perspective images and yielded modest accuracy for 35 classes of moths (43) or used mostly coarse taxonomic resolution (44). We have recently demonstrated that our custom-built time-lapse cameras can record image data from which a deep learning model could accurately estimate local spatial, diurnal, and seasonal dynamics of honey bees and other flower visiting insects (45; Figure 1). Time-lapse cameras are less likely to create observer bias than direct observation and data collection can extend across full diurnal and even seasonal time scales. Cameras can be baited just as traditional light and pheromone traps or placed over ephemeral natural resources such as flowers, fruits, dung, fungi or carrion. Bjerge, *et al*. (46) propose to use an automated light trap to monitor the abundance of moths and other insects attracted to light. The solution is powered by a solar panel, which allows the system to be installed in remote locations (Figure 2). Ultimately, true ‘Internet of Things’ enabled hardware will make it possible to implement classification algorithms directly on the camera units to provide fully autonomous systems in the field to monitor insects and report detection and classification data back to the user or to online portals in real time (34).

**Figure 1.**
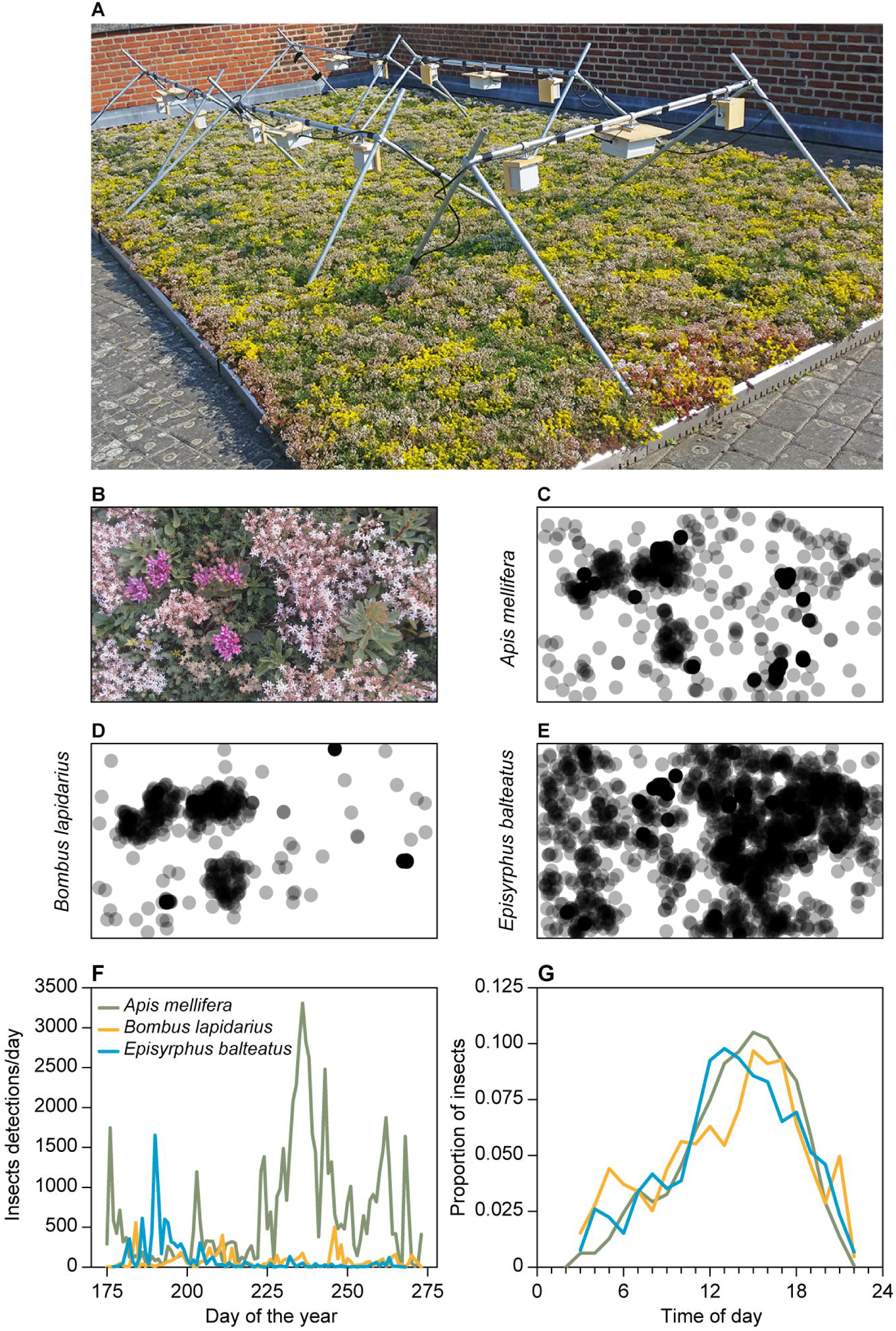
We developed and tested a camera trap for monitoring flower visiting insects, which records images at fixed intervals (45). (A) The setup consist of two web cameras connected to a control unit containing a Raspberry Pi computer and a hard drive. In our test, ten camera traps were mounted on custom built steel rod mounts c. 30cm above a green roof mix of plants in the genus *Sedum*. Images were recorded every 30 sec during the entire flowering season. After training a convolutional neural network (Yolo3), we detected >100,000 instances of pollinators over the course of an entire growing season. (B) An example image from one of the cameras showing a scene consisting of different flowering species. The locations of the insect detections varied greatly among three common flower visiting species (C) the European honey bee (*Apis mellifera)*, (D) the red-tailed bumblebee (*Bombus lapidarius*), and (E) the marmalade hoverfly (*Episyrphus balteatus*). Across the ten camera traps, the deep learning model detected detailed variation in (F) seasonal and (G) diurnal variation in the occurrence frequency among the same three species. Figure adapted with permission from (45).

**Figure 2.**
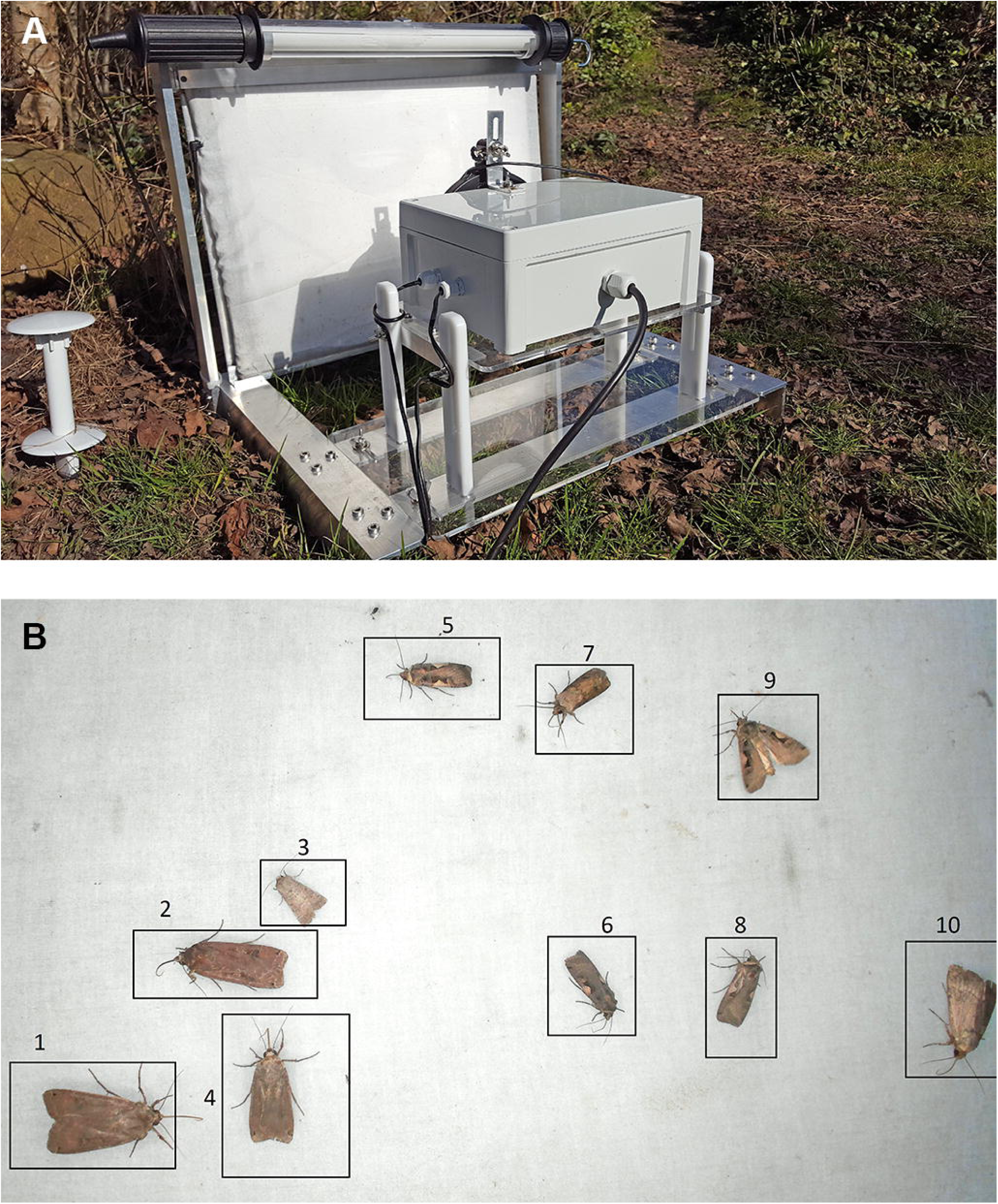
(A) To automatically monitor nocturnal moth species, we designed a light trap with an on-board computer vision system (46). The light trap is equipped with three different light sources. A fluorescent tube to attract moths, a light table covered by a white sheet to provide a diffuse background illumination of the resting insects, and a light ring to illuminate the specimens. The system is able to attract moths and automatically capture images based on motion detection. The trap is designed using standard components such as a high-resolution USB web camera and a Raspberry Pi computer. (B) We have proposed a computer vision algorithm that, during offline processing of the captured images, performs tracking and counting of individual moths. A customized convolutional neural network was trained to detect and classify eight different moth species. The algorithm can run on the on-board computer to allow the system to automatically process and submit species data via a modem to a server. The system works off grid due to a battery and solar panel.

### Radar, acoustic, and other solutions for *in situ* monitoring

The use of radar technology in entomology has allowed for the study of insects at scales not possible with traditional methods, specifically related to both migratory and non-migratory insects flying at high altitudes (47). Utilizing data from established weather radar networks can provide information at the level of continents (48), while specialized radar technology such as vertical-looking radars (VLRs) can provide finer grained data albeit at a local scale (49). The VLRs can give estimates of biomass and body shape of the detected object, and direction of flight, speed and body orientation can be extracted from the return radar signal (50). However, VLR data provide little information on community structure and conclusive species identification requires aerial trapping (51, 52). Harmonic scanning radars can detect insects flying at low altitudes at a range of several hundred meters, but insects need to be tagged with a radar transponder and must be within line-of-sight (53, 54). Collectively, the use of radar technology in entomology can provide valuable information in insect monitoring, for example on the magnitude of biomass flux stemming from insect migrations (55), but requires validation with conventional monitoring methods (e.g. 56).

Bioacoustics is a well-established scientific discipline and acoustic signals have been extensively and widely used in the field of ecology, for example for detecting presence and studying behavior of marine mammals (57) and for bird species identification (58). Jeliazkov, *et al*. (59) used audio recordings to study population trends of Orthoptera at large spatial and temporal scales, demonstrating that bioacoustic techniques have merit in entomological monitoring. Machine learning methods have proven a particularly valuable tool for deciphering noisy audio recordings and detecting the signals of animals. Kiskin, *et al*. (60) demonstrated the use of a CNN to detect the presence of mosquitoes by identifying the acoustic signal of their wingbeats. Other studies have shown that even species classification can be done using machine learning on audio data, for example for birds (58), bats (61), grasshoppers (62), and bees (63). Although, it has been argued that the use of pseudo-acoustic optical sensors rather than actual acoustic sensors is a more promising technology because of the much improved signal-to-noise ratio in these systems, which may be a particularly important point for bioacoustics in entomology (64).

Other systems rely on sensor technology to automate the recording of insect activity or even body mass, but without actual consideration of the subsequent processing of the data with deep learning methods (65, 66). In (65) they use a sensor-ring of photodiodes and infrared LEDs to detect large and small sized arthropods, including pollinators and pests and achieve a 95% detection accuracy for live microarthropods of three different species in the size range of 0.5 – 1.1 mm. The Edapholog (66) is a low-power monitoring system for real-time detection of soil microarthropods where a pitfall trap is presented. Probe and sensing is based on detection of change in infrared light intensity similar to (65) and it counts the organisms falling into the trap and estimates their body size. The probe is connected via radio signals to a logging devices that transmits the data to a central server for real-time monitoring. Similarly, others have augmented a traditional low-cost trapping methods by implementing optoelectronic sensors and wireless communication to allow for real-time monitoring and reporting (35). Since, such sensors do not produce images that are intuitive to validate, it could be challenging to generate sufficient, validated training data for implementing deep learning models, although such models could still prove useful.

### Digitizing specimens and natural history collections

There are strong efforts to digitize natural history collections for multiple reasons including the benefits of deep learning applications (67). The need for and benefits of digitizing natural science collections have motivated the foundation of the Distributed System of Scientific Collections Research Infrastructure (DISSCo RI, www.dissco.eu). DISSCo RI strives for the digital unification of all European natural science assets under common curation and access policies and practices. Most existing databases include single view digitisations of pinned specimens (68), while datasets of insect specimens recorded using multiple sensors, 3D models, and databases on living insect specimens are only just emerging (69, 70). The latter could be particularly relevant for deep learning models. There is also a valuable archive of entomological data in herbarium specimens in the form of signs of herbivory (71). The standard digitization of herbarium collections has proven suitable for extracting herbivory data using machine learning techniques (72). Techniques to automate digitization techniques will accelerate the development of such valuable databases and enables tools for identification of non-pinned specimens and live insects *in situ* (67). The BIODISCOVER machine (73) is a proposal towards the automatization of creating databases of liquid preserved specimens such as most field collected insects. The process consists of four automatized steps: 1) bin picking of individual insects directly from bulk samples, 2) recording the specimen from multiple angles using high speed imaging, 3) saving the captured data in an optimized way for deep learning algorithm training and further study, and 4) sorting specimens according to size, taxonomic identity or rarity for potential further molecular processing (Figure 3). Digitization efforts should carefully consider how image data of specimens can be leveraged in efforts to develop deep learning models for *in situ* monitoring.

**Figure 3.**
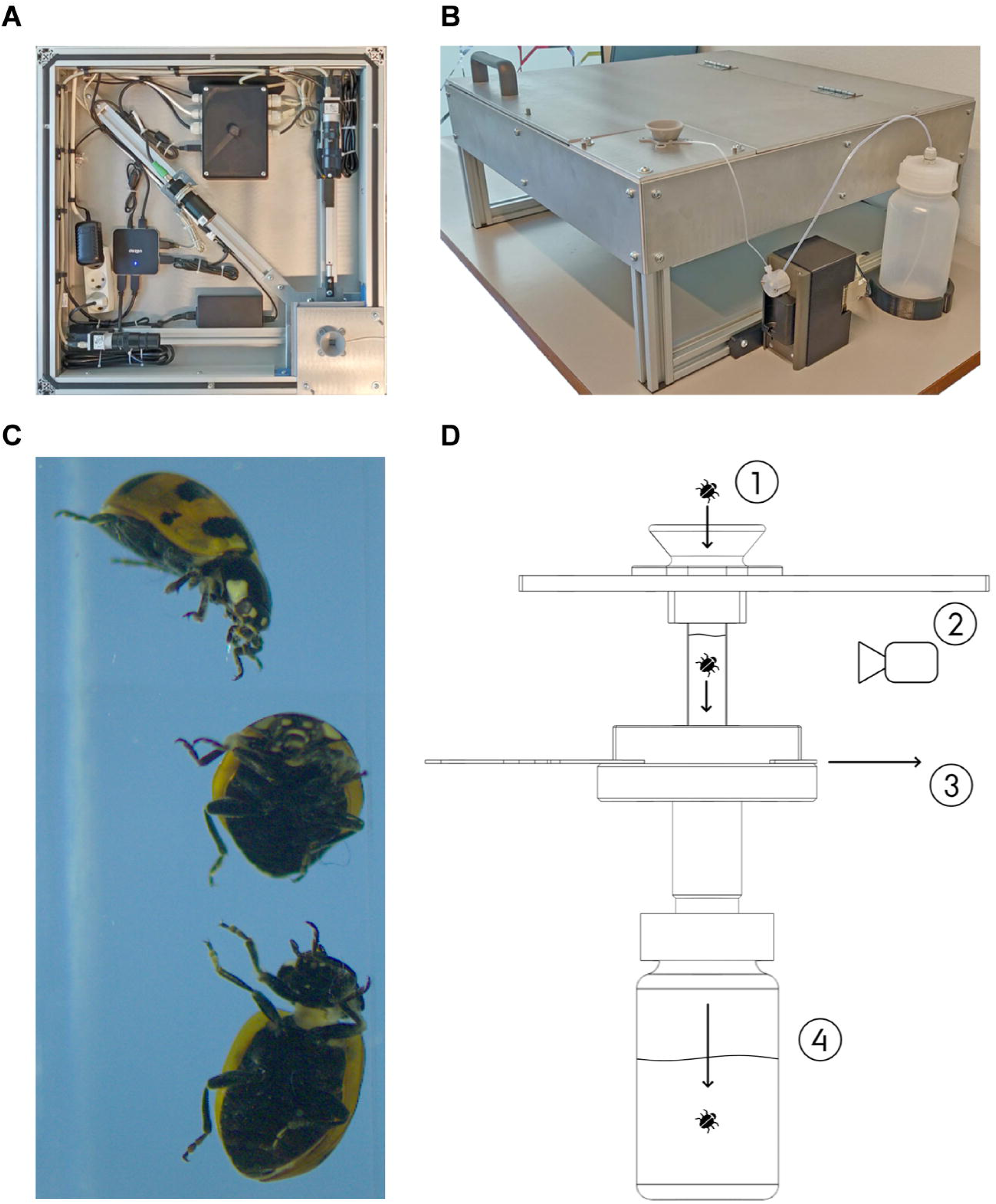
The BIODISCOVER machine, can automate the process of invertebrate sample sorting, species identification, and biomass estimation (73). (A) The imaging system consists of an ethanol-filled spectroscopic cuvette, a powerful and adjustable light source and two cameras capable of recording images at 50 frames per second (B) The setup is mounted in a light proof aluminium box and fitted with a pump for refilling the spectroscopic cuvette. (C) Each specimen is imaged from two angles by the cameras as it is dropped into the ethanol-filled cuvette and geometric features related to size and biomass are computed automatically. (D) The system has a built in flushing mechanism for controlling which specimens should be kept together for subsequent storage or analysis. The results for an initial dataset of images of 598 specimens across 12 species of known identity was very promising with a classification accuracy of 98.0% (73). The system is generic and can easily be used for other groups of invertebrates as well. As such, the BIODISCOVER machine pave the way for cheap, fast, and accurate data on spatial and temporal variation in invertebrate abundance, diversity and biomass. Figure adapted with permission from (73).

## POTENTIAL DEEP LEARNING APPLICATIONS IN ENTOMOLOGY

The big data collected by sensor-based insect monitoring as described above requires efficient solutions for transforming the data into biologically relevant information. Preliminary results suggest that deep learning offers a valuable tool in this respect and could further inspire the collection of new types of data (20, 45). Deep learning software, e.g. for ecological applications, is mostly constructed using open source Python libraries and frameworks such as TensorFlow, Keras, PyTorch, and Scikit-learn (24) and prototype implementations are typically publicly available e.g. on www.github.com. This, in turn, makes the latest advances in other fields related to object detection and fine-grained classification available also for entomological research. As such, the existing deep learning toolbox is already available, but will need adaptation to entomology from the domains for which the tools were developed. In the following, we provide a brief description of the transformative potential of deep learning related to entomological data stored in images structured around four main applications.

### Detecting and tracking individuals *in situ*

Image-based monitoring of insect abundance and diversity could rapidly become globally widespread as countries make efforts to better understand the severity of the global insect decline and mitigation measures. Similarly, tracking of individual insects *in situ* even for short periods of time holds exciting research potential. For example, by estimating movement speed of individual insects in their natural environments and relating it to observed microclimatic variation, more realistic thermal performance curves can be established and contrasted to traditional lab-derived thermal performance. However, tracking insect in their natural environment is currently a highly challenging task, due to e.g. the cluttered scenes and varying lighting conditions. In computer vision, such tasks are termed ‘detection-based online multiple object tracking’, and work under a set of assumptions (74). These assumptions include a precise initial detection (initialization) of the objects to be tracked in a scene, a good ability to visually discriminate between the multiple tracked objects, and smooth motion, velocity, and acceleration patterns of the tracked objects (75). The small visual differences among individual insects and frequent hiding behaviour violate the above assumptions. Moreover, current state-of-the-art deep learning models typically use millions of learned parameters and can only run in near real-time with low-resolution video, which constrains the visual discrimination of the targeted objects in the scene. Possible solutions to these challenges include the use of non-linear motion models (76) and the development of compact (77) or compressed (78) deep learning models.

If we manage to solve the task of individual tracking of insects it could open the doors for a new individual-based ecology with profound impacts in such research fields as population, behavioural, and thermal ecology as well as conservation biology. Moreover, considering the recent development in low-cost powerful graphical processing units and dedicated artificial intelligence processer suitable for autonomous and embedded systems (e.g. NVIDIA Jetson Nano, Google Coral Edge TPU, and the Intel AI USB stick), it may soon become feasible to detect, track, and decode behaviour of insects in real-time and report information back to the user.

### Detecting species interactions

Species interactions are critical for the functioning of ecosystems, yet as they are ephemeral and fast, the consequences of a disruption for ecological function is hard to quantify. High temporal resolution image-based monitoring of consumers and resources can allow for a unique quantification of species interactions (79). The use of cameras allows for continuous observations of species and their interactions across entire growing seasons such as insects visiting flowers, defoliation by herbivores, and predation events. There is an urgent need to develop methods to observe and quantify species interactions efficiently and at ecologically relevant spatial and temporal scales (80, 81). To detect such interactions, image recording should be collected at the scales where individuals interact, i.e., by observing interacting individuals at intervals of seconds to minutes, yet they should ideally extend over seasonal and/or multi-annual periods, which at the moment is difficult to fulfil. Our preliminary results have demonstrated an exciting potential to record plant-insect interactions using time-lapse cameras and deep learning (28 and Figure 1).

### Taxonomic identification

Taxonomic identification can be approached as a deep learning classification problem. Deep learning-based classification accuracies for image-based insect identification of specimens are approaching the accuracy of human experts (82-84). Applications of gradient-weighted class activation mapping can even visualize morphologically important features for CNN classification (84). Classification accuracy is generally much lower when the insects are recorded live in their natural environments (85, 86), but when class confidence is low at the species-level, it may still be possible to confidently classify insects to a coarser taxonomic resolution (87). In recent years, impressive results have been obtained by CNNs (88). They can classify huge image datasets, such as the 1000-class ImageNet dataset at high accuracy and speed (89). With images of >10,000 species of plants, classification performance of CNNs is currently much lower than for botanical experts (25), but promising results in distributed training of deep neural networks (90) and federated learning (91, 92) suggest that improvements can be expected.

In most ecological communities, it is common for species to be rare. This often results in highly imbalanced datasets, and the number of specimens representing the rarest species could be insufficient for training neural networks (86, 87). As such, advancing the development of algorithms and approaches for improved identification of rare classes is a key challenge for deep learning-based taxonomic identification. Solutions to this challenge could be inspired by class resampling and cost-sensitive training (93) or by multiset feature learning (94, 95). Class resampling aims at balancing the classes by under-sampling the larger classes and/or over-sampling the smaller classes, while cost-sensitive training assigns a higher loss for errors on the smaller classes. In multiset feature learning, the larger classes are split into smaller subsets, which are combined with the smaller classes to form separate training sets. These methods are all used to learn features that can more robustly distinguish the smaller classes. Species identification performance can vary widely, ranging from species which are correctly identified in most cases to species that are generally difficult to identify (96). Typically, the amount of training data is a key element for successful identification, although recent analyses of images of the approximately 65,000 specimens in the carabid beetle collection at the Natural History Museum London suggest that imbalances in identification performance are not necessarily related to how well-represented a species is in the training data (87). Further work is needed on large datasets to fully understand these challenges.

A related challenge is formed by those species that are completely absent from the reference database on which the deep learning models are trained. Detecting such species requires techniques developed for multiple-class novelty/anomaly detection or open set/world recognition (97, 98). A recent survey introduces various open set recognition methods with the two main approaches being discriminative and generative (99). Discriminative models are based on traditional machine learning techniques or deep neural networks with some additional mechanism to detect outliers, while the main idea of generative models is to generate either positive or negative samples for training. However, the current methods are typically applied on relatively small datasets and do not scale well with the number of classes (99). Insect datasets typically have a high number of classes and a very fine-grained distribution, where the phenotypic differences between species may be minute while intra-species differences may be large. Such datasets are especially challenging for open set recognition methods. While it will be extremely difficult to overcome this challenge for all species using only phenotype based identification, combining image-based deep learning and DNA barcoding techniques may help to solve the problem.

### Estimating biomass from bulk samples

Deep learning models can potentially predict biomass of bulk insect samples in a lab setting. Legislative aquatic monitoring efforts in the United States and Europe require information about the abundance or biomass of individual taxa from bulk invertebrate samples. Using the BIODISCOVER machine, Ärje, *et al*. (73) were able to estimate biomass variation of individual specimens of Diptera species without destroying specimens. This was achieved from geometric features of the specimen such as the mean area from multiple images recorded by the BIODISCOVER machine and statistically relating such values to subsequently obtained dry mass from the same specimens. To validate such approaches, it is necessary to have accurate information about the dry mass of a large selection of taxa. In the future, deep learning models may provide even more accurate estimates of biomass. Obtaining specimen-specific biomass information non-destructively from bulk samples is a high priority in routine insect monitoring, since it will enable more extensive insights into insect population and community dynamics and provide better information for environmental management.

## FUTURE DIRECTIONS

To unlock the full potential of deep learning methods for insect ecology and monitoring, four main challenges need to be addressed with highest priority. We describe each of these below.

### Validating image-based taxonomic identification

Validation of the detection and identification of species recorded with cameras in the field pose a critical challenge for implementing deep learning tools in entomology. Often it will not be possible to conclusively identify insects from images and validation of image-based species classification should be done using alternative, complimentary techniques. We suggest four approaches to this validation: 1) Obtaining local knowledge about the identity and relative abundance of candidate species, 2) catching and manually identifying insects in the vicinity of a camera trap, 3) identifying insects by environmental DNA analysis of insect DNA traces left e.g. on flowers (100), or 4) by directly observing and catching insects visible to the camera. The first three approaches are indirect and each come with their separate problems such as the difference in trapping efficiency of a time-lapse camera trap and e.g. a pitfall trap placed to capture the same insects. However, the subsequent identification of specimens from pitfall trapping can serve as validation of image-based results and can further help in production of training data for optimizing deep learning models (e.g. by placing specimens back under the camera). DNA techniques be able to validate image-based identification since DNA can give accurate information on species identity (11, 100, 101).

For specific purposes, validation of insects can be done through interfaces with online portals and by involving citizen science. With integrated deep learning algorithms, online portals provide instant candidate species when users upload pictures of observed insect species. The most prominent examples of such portals of relevance to insects are the smartphone apps connected to sites such as www.iNaturalist.org and www.observation.org. Another way of using deep learning models to generate data on insect occurrence in their natural environment is by involving the public in the annotation and quality control of images of insects uploaded to citizen science web portals such as www.zooniverse.org (102).

### Generating training data

One of the main challenges with deep learning is the need for large amounts of training data, which is slow, difficult, and expensive to collect and label. Deep learning models typically require hundreds of training instances of a given species to learn to detect species occurrences against the background (86). In a laboratory setting, the collection of data can be eased by automated imaging devices, such BIODISCOVER described above, which allow imaging large amounts of insects under fixed settings. The imaging of species *in situ* should be done in a wide range of conditions (e.g., background, time of day, and season) to avoid that the model learns a false connection between the species and the background, with resulting lower ability of the model to detect the species against another background. Approaches to alleviate the challenge of moving from one environment to another include multi-task learning (103), style transfer (104), image generation (105), or domain adaptation (106). Multi-task learning aims to concurrently learn multiple different tasks (e.g., segmentation, classification, detection) by sharing information leading to better data representations and ultimately better results. Style transfer methods try to impose properties appearing in one set of data to new data. Image generation can be used to created synthetic training images with, for example, varying backgrounds. Domain adaptation aims at tuning the parameters of a deep learning model trained on data following one distribution (source domain) to adapt so that they can provide high performance on new data following another distribution (target domain).

The motion detection sensors in wildlife cameras are typically not triggered by insects and species typically only occur in a small fraction of time-lapse images. A key challenge is therefore to detect insects and filter out blank images from images with species of interest (102, 107). When it is difficult to obtain sufficient samples of rare insects, Zhong, *et al*. (108) proposed to use deep learning only to detect all species of flying insects as a single class. Subsequently, the fine-grained species classification can be based on manual feature extraction and support vector machines, which is a machine learning technique that requires less training data and solves the problem of insufficient training data.

The issue of scarce training data can also be alleviated with new data synthesis. Data synthesis could be used specifically to augment the training set by creating artificial images of segmented individual insects that are placed randomly in scenes with different backgrounds (109). A promising alternative is to use deep learning models for generating artificial images belonging to the class of interest. The most widely approach to date is based on generative adversarial networks (110) and has shown astonishing performance results in computer vision problems in general, as well as in ecological problems (111).

### Building reference databases

Publicly available reference databases are critical for adapting deep learning tools to entomological research. Initiatives like DISSCO RI and IDigBio (https:www.idigbio.org/) are important for enabling the use of museum collections. However, to enable deep learning-based identification, individual open datasets from entomological research and monitoring are also needed (e.g. 85, 96, 112). The collation of such research datasets will require dedicated projects as well as large coordinated efforts that drive the open-access and reuse of research data such as the European Open Science Cloud and the Research Data Alliance. Building a large insect reference dataset is laborious and, therefore, it is important to maximize the benefits. To do so, non-collection datasets should also use common approaches and hardware and abide to best practices in metadata and data management (113-115). Further, dataset collectors and deep learning model developers should work closely together and make data accessible. All the possible metadata, such as camera settings and hardware, sampling location, date, and time of day, should be saved for future analysis. Similarly, characteristics of the specimen, such as species identity, biomass, sex, age class, and possibly derived information like dry weight should be recorded if such information exist. In particular, correct labelling of species in images is critical. Using multiple experts and molecular information about species identity to verify the labeling or performing subsequent validity checks through DNA barcoding will improve the data quality and the performance of the deep learning models. This can be done, for instance, by manually verifying the quality and labeling of images that are repeatedly misclassified by the machine learning methods. Standardized imaging devices such as the BIODISCOVER machine could also play a key role in building reference databases from monitoring programs (73). Training classifiers with species that are currently not encountered in a certain region but can possibly spread there later will naturally help to detect such changes when they occur. Integration of such reference databases with field monitoring methods forms an important future challenge. As a starting point, we provide a list of open access entomological image databases (SI Appendix).

### Integration of deep learning and DNA-based tools

For processing samples in the lab, molecular methods have gained increasing attention over the past decade, but there are still critical challenges which remain unresolved: specimens are typically destroyed, abundance cannot be accurately estimated, and key specimens cannot be identified in bulk samples. Nevertheless, DNA barcoding is now an established, powerful method to reliably assess biodiversity also in entomology (11). For insects, this works by sequencing a short fragment of the mitochondrial cytochrome-c-oxidase I subunit gene (COI) and comparing the DNA sequence to an available reference database (116). Even undescribed and morphologically cryptic species can be distinguished with this approach (117), which is unlikely to be possible with deep learning. This is of great importance as morphologically similar species can have distinct ecological preferences (118) and thus distinguishing them unambiguously is important for monitoring, ecosystem assessment and conservation biology. However, mass-sequencing based molecular methods cannot provide precise abundance or biomass estimates and assign sequences to individual specimens (12). Therefore, an unparalleled strength lies in combining both image-recognition and DNA metabarcoding approaches: i) When building reference collections for training models for insect classification, species identity can be molecularly verified and potential cryptic species can be separated by the DNA barcode. ii) After image-based species identification of a whole bulk sample, all specimens can be processed via DNA metabarcoding to assess taxonomic resolution at the highest level. A further obvious advantage of linking computer vision and deep learning to DNA is the fact that even in the absence of formal species descriptions, DNA tools can generate distinctly referenced taxonomic assignments via so-called “Barcode-Index-Numbers” (BINs) (119). These BINs provide referenced biodiversity units using the taxonomic backbone of the Barcode of Life Data Systems (https://boldsystems.org) and represent a much greater diversity of even yet undescribed species. For instance, it is typically clear that a new species belongs to the genus *Astraptes* in the butterfly family Hesperiidae, but also that it represents a genetically distinct, new entity (120). These units can also be directly used as part of ecosystem status assessment despite not yet having Linnean names. BINs can be used for model training. Recent studies convincingly show that with this more holistic approach, which includes cryptic and undescribed species, the predictions of environmental status as required by several legislative monitoring programs actually improve substantially (e.g. 121). For cases of cryptic species with great relevance e.g. for conservation biology it is also possible to individually process specimens of a cryptic species complex after automated image-based assignment to further validate identity and frequency of these. Combining deep learning with DNA-based approaches could deliver detailed trait information, biomass, and abundance with the best possible taxonomic resolution.

## CONCLUSION

Deep learning is currently influencing a wide range of scientific disciplines (88), but has only just begun to benefit entomology. While there is a vast potential for deep learning to transform insect ecology and monitoring, applying deep learning to entomological research questions brings new technical challenges. The complexity of deep learning models and the challenges of entomological data require substantial investment in interdisciplinary efforts to unleash the potential of deep learning in entomology. However, these challenges also represent ample potential for cross-fertilization among the biological and computer sciences. The benefit to entomology is not only more data, but also novel kinds of data. As the deep learning tools become widely available and intuitive to use, they can transform field entomology by providing information that is currently intractable to record by human observations (18, 33, 122). Consequently, there is a bright future for entomology, with new research niches opening up and access to unforeseen scales and resolution of data, vital for biodiversity assessments.

The shift towards automated methods may raise concerns about the future for taxonomists, much like the debate concerned with developments in molecular species identification (123, 124). We emphasize that the expertise of taxonomists is at the heart of and critical to these developments. Initially, automated techniques will be used in the most routine-like tasks, which in turn will allow the taxonomic experts to dedicate their focus on the specimens requiring more in depth studies as well as the plethora of new species that need to be described and studied. To enable this, we need to consider approaches that can pinpoint samples for human expert inspection in a meaningful way, e.g., based on neural network classification confidences (82) or additional rare species detectors (125). As deep learning becomes more closely integrated in entomological research, the vision of real-time detection, tracking, and decoding of behaviour of insects could be realized for a transformation of insect ecology and monitoring. In turn, efficient tracking of insect biodiversity trends will aid the design of effective measures to counteract or revert biodiversity loss.

## ACKNOWLEDGEMENTS

David Wagner is gratefully thanked for convening the session “Insect declines in the Anthropocene” at the Entomological Society of America annual meeting 2019 in St. Louis, USA, which brought the group of contributors to the special feature together. TTH acknowledges funding from the Villum Foundation (grant 17523) and the Independent Research Fund Denmark (grant 8021-00423B), Kristian Meissner acknowledges funding from the Nordic Council of Ministers (project 18103, SCANDNAnet). Jenni Raitoharju acknowledges funding from Academy of Finland (project 324475).

---

TABLE 1

## Glossary

Bin picking: an industrial term for robots that pick up one of many objects randomly placed in a container.
Convolutional Neural Network (CNN): a deep learning algorithm in the family of neural networks with serval different layers commonly applied for image recognition and classification. A CNN can be trained to recognize various objects and patterns in an image. There are four main different operations in a CNN: convolution, activation functions, sub sampling, and fully connected layer. During training the learnable parameters of each convolutional and fully connected layer are adjusted so the CNN is able to recognize different patterns of the training data and used for final image classification.
Data augmentation: a technique that can be used to artificially expand the size of a training dataset by creating modified images with objects of interest for classification.
Machine learning: a subset of artificial intelligence associated with creating algorithms that can change themselves without human intervention to get the desired result – by feeding themselves through structured data.
Deep learning: a subset of machine learning where algorithms are created and function similarly to machine learning, but where there are many levels of these algorithms, each providing a different interpretation of the data it conveys.
DNA barcoding: Identification of a species using a short, standardised gene fragment.
Initialization: description of an object to be tracked.
Training data: classified images (e.g. images of known species identified by experts) that are recorded to train a deep learning model.
Precision: the number of true positives divided by the sum of true positives and false positives
Recall: also called the true positive rate, is the number of true positives divided by the sum of true positives and false negatives.
Classification accuracy: the sum of true positives and true negatives divided by the total number of specimens.

